# A empirical comparison of preservation methods for synthetic DNA data storage

**DOI:** 10.1101/2020.09.19.304014

**Authors:** Lee Organick, Bichlien H. Nguyen, Rachel McAmis, Weida D. Chen, A. Xavier Kohll, Siena Dumas Ang, Robert N. Grass, Luis Ceze, Karin Strauss

## Abstract

Synthetic DNA has recently risen as a viable alternative for long-term digital data storage. To ensure that information is safely recovered after storage, it is essential to appropriately preserve the physical DNA molecules encoding the data. While preservation of biological DNA has been studied previously, synthetic DNA differs in that it is typically much shorter in length, it has different sequence profiles with fewer, if any, repeats (or homopolymers), and it has different contaminants. In this paper we evaluate nine different methods used to preserve data files encoded in synthetic DNA by accelerated aging of nearly 29,000 DNA sequences. In addition to a molecular count comparison, we also sequence and analyze the DNA after aging. Our findings show that errors and erasures are stochastic and show no practical distribution difference between preservation methods. Finally, we compare the physical density of these methods and provide a stability versus density trade-offs discussion.

## Introduction

Synthetic DNA has been growing in popularity as a promising new technology for long-term digital data storage^1–4^. DNA is very dense, with expected practical densities higher than one exabyte (10^18^ bytes) per cubic inch. This is orders of magnitude higher than current storage media. Unlike other media, whose reading technology quickly becomes obsolete as the technology evolves, DNA is expected to always be readable and compatible with existing and future DNA sequencing platforms due to its prominence in life sciences and clinical applications. Long-term digital data storage relies on the integrity of the physical medium for data endurance over decades up to possibly thousands of years. This is no exception for digital data storage in synthetic DNA. After storage, anywhere from 10 to over 1000 intact copies of each sequence are required for data retrieval^2, 5^.

Biological DNA samples are commonly frozen or chilled significantly below room temperature for preservation. The simplicity of this method makes it attractive in a laboratory setting for storing small quantities of DNA. However, this is significantly less appealing for DNA data storage due to a different set of constraints to reach wide deployment: access to the DNA must be fully automated, density must be kept as high as possible, and the cost and energy to store and maintain the samples as low as possible.

For these reasons, there has been growing interest at storing DNA at room temperature. Most notably, individual studies examine synthetic DNA preservation at room temperature with nanoparticles, encapsulation in metal capsules, trehalose, and various other sugar matrices^6, 7^. While there are a number of commercial products designed to preserve DNA, there have been no large, comprehensive comparison studies of the stability of digital data storage in synthetic DNA under the various methods until now. We selected DNA preservation methods for evaluation by factoring in protocol simplicity and reported ability to store DNA at room temperature. The most “primitive” methods chosen were to store DNA dehydrated in a standard polypropylene eppendorf tube (referred to as “No Additives”) and to store DNA dehydrated on filter paper, a relatively old method of preserving and transporting biological specimens dating back to the 1960s^8^. The next set of methods involved mixing DNA with various additives prior to dehydration in an eppendorf tube. These methods included using trehalose (referred to as “Trehalose”), a disaccharide that enables several organisms to survive desiccation^9^, and also a mixture of sugars comprised of trehalose, raffinose, maninitol, and uric acid (“Sugar Mix”). Similarly, we included commercially available proprietary sugar mixtures advertised for room temperature DNA storage (“DNAStable” and “GenTegra”). We also investigated more sophisticated encapsulation techniques to preserve DNA. These methods included storing DNA in Imagene DNAshells (“Imagene”), in which DNA is stored in a borosilicate glass insert inside a stainless steel shell and cap, which is hermetically sealed and filled with non-reactive gas, and on magnetic nanoparticles further coated with DNAStable (“Mag-Bind DNAStable”). While all of these preservation methods were sequenced immediately after rehydration or de-encapsulation with no further preparation or manipulation, we also examined our ability to manipulate a pool of DNA after preservation with a preparation method that included polymerase chain reaction, PCR, (“DNAStable + PCR”).

We present an analysis of these different DNA storage methods used to preserve synthetic DNA encoding two different data files. We performed accelerated aging of the samples at 65°*C*, 75°*C*, and 85°*C* to determine the first order decay kinetics of each storage method and analyzed the sequencing data to determine the percentage of sequences recovered. We also tested DNA preservation in four methods at a different lab (ETH-Z) in addition to those performed at our UW lab. Three preservation methods were tested in both labs “No Additives”, “Trehalose”, “DNAStable”) and one was performed solely at ETH-Z (“Magnetic NP”).

## Results

### qPCR of DNA Material

To evaluate the effectiveness of each storage method, we selected a small data file encoded in 7,373 unique DNA sequences and a larger data file encoded in 21,601 unique sequences. The overall architecture of these sequences features a 110bp payload encoding the digital data flanked by a 20bp forward primer and a 20bp reverse primer, resulting in a sequence 150bp long in total. Each digital file has a unique pair of forward and reverse primer sequences (the details of this architecture are reported in previous work^4^).

The two files were amplified individually via PCR and combined into one solution, totalling 28,974 unique sequences. Aliquots of this solution were used for the aging experiment. Triplicates of each aliquot were subjected to accelerated aging at three storage temperatures (65^°^*C*, 75^°^*C*, 85^°^*C*) for five different time points. All samples were kept in ovens maintaining 50% relative humidity. More details about each storage method can be found in their respective Methods sections. To reduce errors associated with sequencing preparation, each sample was ligated with Illumina adapters and tagged with an unique index prior to the accelerated aging experiment and were ready for sequencing directly after aging (note that the “DNAStable + PCR” method was the exception to this, details in Methods). An overview of this experiment is shown in **Fig. 1**.

**Figure 1.**
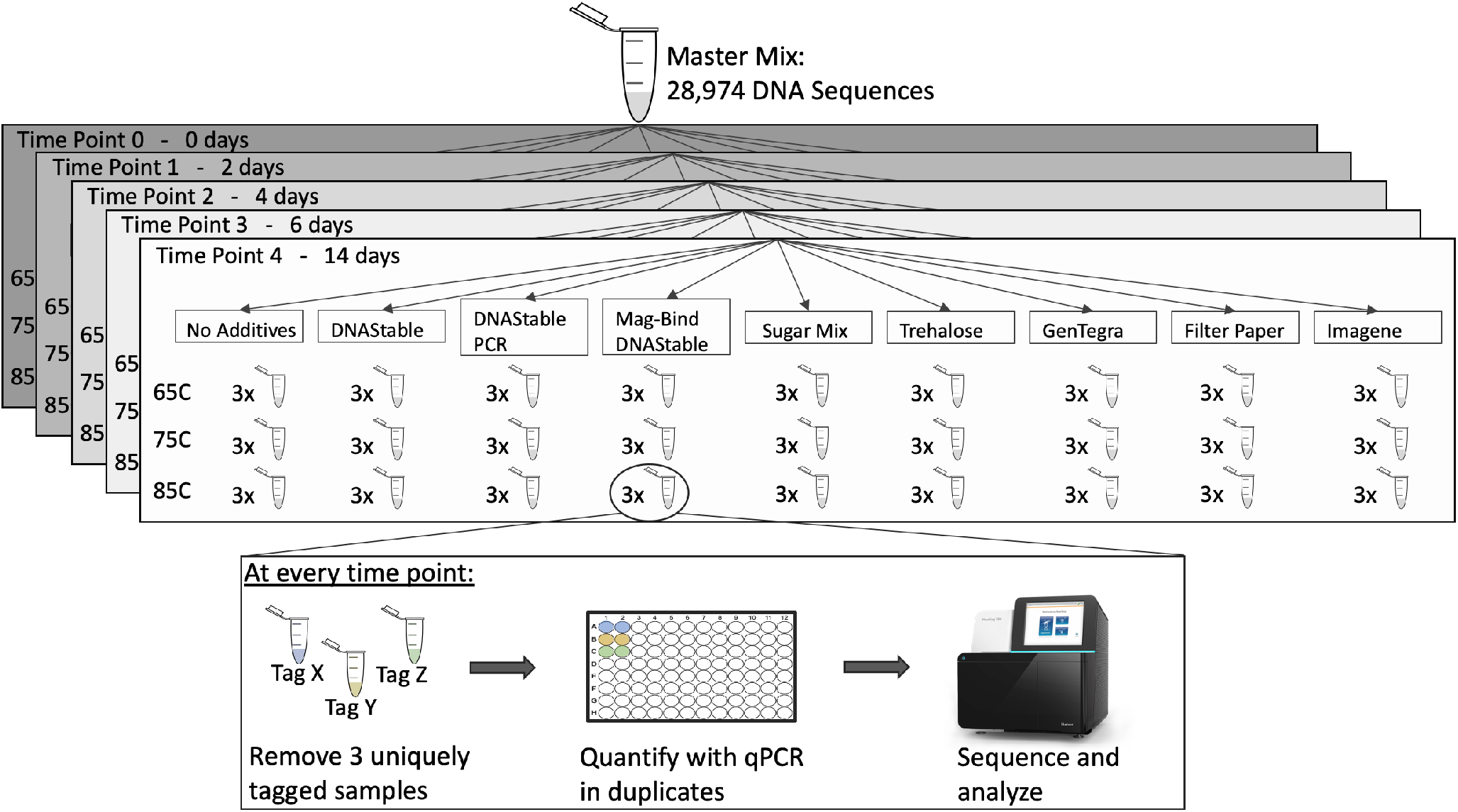
An overview of the aging process. DNA sequences were amplified with PCR and mixed together into one master mix tube. Aliquots were taken from that one tube and preserved with nine different techniques. Each technique and each temperature had three replicates per time point, each replicate with a unique molecular tag (Tag A, Tag B…Tag Z, etc.). At each time point, triplicates were removed from each condition and quantified with qPCR in duplicate, then sequenced with Next-Generation Sequencing (NGS) if there was sufficient material. Note that the time points were different for Imagene and samples aged at ETH-Z as shown in **Fig. 2** and **Fig. 3**, respectively.

At each time point, the concentration of each of the samples was measured by quantitative polymerase chain reaction (qPCR), as shown in **Fig. 2**. Only strands that didn’t break are detectable with qPCR, for exponential amplification is only possible when both the forward and reverse primer are present on the same DNA strand. While all DNA preservation methods degraded at a slower rate than only dehydrating the sample (No Additives), there was no clear ranking of preservation methods only looking at one experimental temperature. For example, GenTegra consistently degraded slower than Trehalose at 65°*C*, but not at 75°*C*. This could be a result of the various proprietary ingredients included in the GenTegra mix having different temperature-dependent protective properties. The GenTegra user guide specifies that the GenTegra DNA preservation material is “designed to tolerate temperatures of −80C to 76C during transport”^13^.

**Figure 2.**
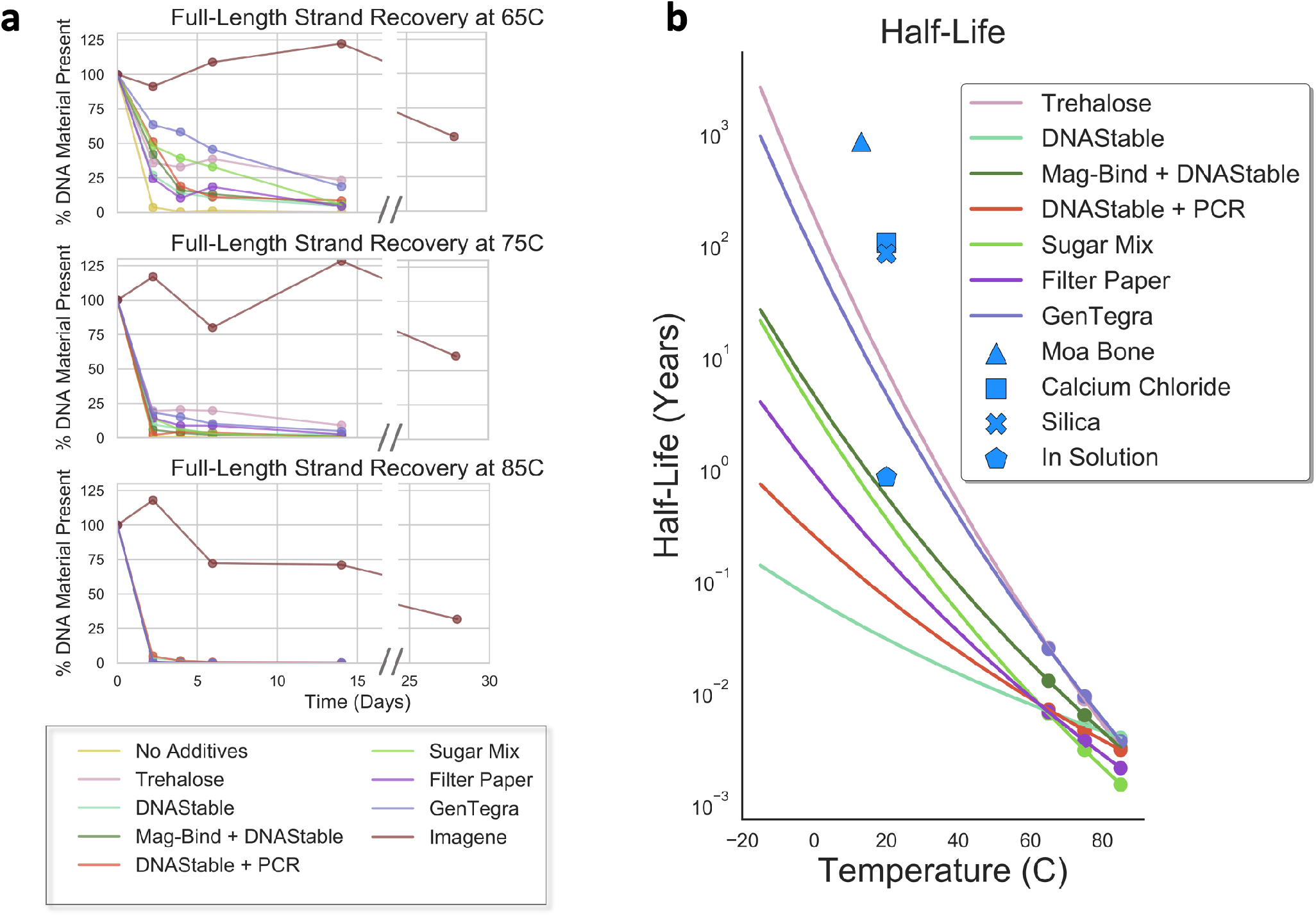
DNA degradation results. (a) The percent of full-length DNA material present after each time point, as measured with a quantitative polymerase chain reaction (qPCR). (b) Extrapolated DNA half life of 150bp DNA strands at various temperatures for all preservation methods that had measurable qPCR data for at least two time points for each temperature. Our Imagene data is not shown because it did not degrade enough to reliably measure a rate of decay, however it has been extrapolated in prior work^10^ and is shown here in blue (“DNAshell”). The five data points in blue are shown as a comparison to other reported DNA preservation results in literature (DNAshell technology from Imagene^10^, Moa bone^11^, calcium chloride^6^, silica^7^, in solution^12^). All data are scaled to reflect the half life of a 150bp DNA strand following previously established scaling methods^7^ of t_1/2_^150bp^ = t_1/2_^1bp^/150. For more calculation information, see **Supplemental Section 6**.

To compare the various preservation methods more comprehensively, we measured the decay kinetics by incorporating data from all three temperatures and time points. Assuming first-order kinetics, consistent with prior work^7^, we calculated the temperature dependence of the per-nucleotide fragmentation rate (k) using the Arrenhius equation and solved the half-life of the DNA samples at 65°*C*, 75°*C*, and 85°*C* (**Fig. 2**). Details can be found in the Methods section. We did not solve for the half-life of DNA preservation methods that degraded completely by the first time point (No Additives), or did not degrade enough for analysis (Imagene). Our finding that Imagene’s DNAshells degraded exceptionally slowly is consistent with previous work performed with biological samples^13,14^.

Each method preserved DNA better than no preservation material at all (No Additives). Based on the extrapolated half-lives, we found that samples preserved with trehalose and GenTegra had nearly indistinguishable longest half-lives, while the two methods utilizing only DNAStable had the shortest half-lives. However, the method utilizing DNAStable in conjunction with magnetic beads performed nearly identically to samples preserved in the mix of sugars and these two methods had the next shortest half-lives. The filter paper sample degraded only slightly faster than the DNAStable and magnetic beads method and the sugar mix. However, we caution that these extrapolations are sensitive to the amount of DNA material stored and the amount of preservation material, therefore extrapolations shown here are more likely a relative ranking than an exact half-life (see the Discussion section for more details).

### qPCR of ETH-Z Material

Since the samples aged at ETH-Z contained the DNA sequences prior to library preparation, ETH-Z samples were less than half the length of the remaining samples (150 bp and 310 bp, respectively). Therefore, we derived the per-nucleotide fragmentation rate (k) of all samples and scaled all data to the half-life of a 150bp sequence to provide a direct comparison between the two studies, as shown in **Fig. 2** and **Fig. 3**. Note that the preservation methods “No Additives” and “DNAStable” both degraded almost entirely before the first time point at all three temperatures tested and were therefore excluded from the half-life extrapolation.

**Figure 3.**
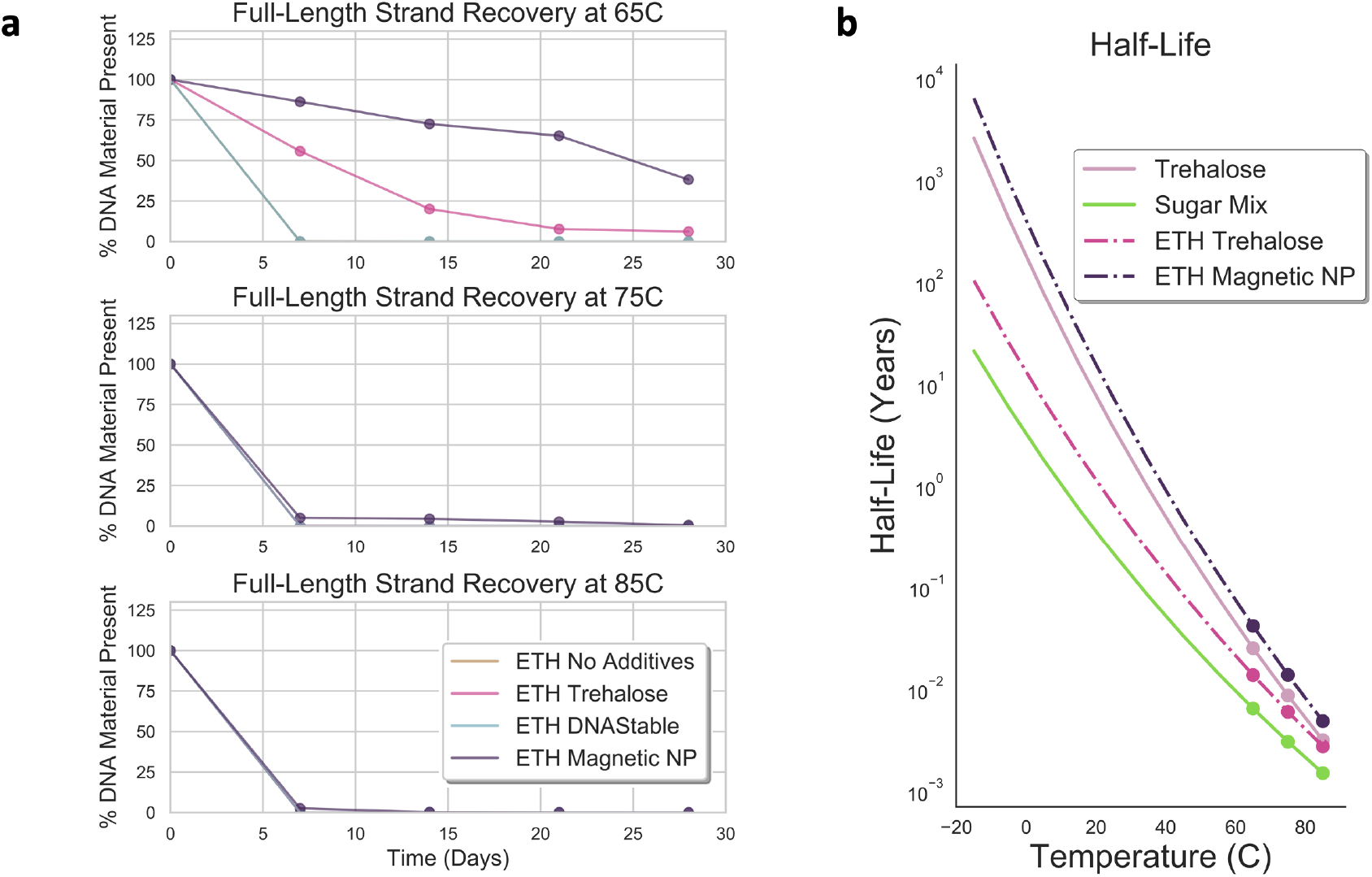
DNA degradation results from ETH-Z and UW. (a) The percent of full-length DNA material present after each time point, as measured with qPCR. (b) Extrapolated DNA half-life at various temperatures for all preservation methods that had measurable qPCR data for at least two time points for each temperature. All data are scaled to reflect the half life of a 150bp DNA strand following previously established scaling methods^7^ of t_1/2_^150bp^ = t_1/2_^1bp^/150. For more calculation information, see **Supplemental Section 6**.

We observed a significant difference in decay kinetics between the UW and ETH-Z samples stored with trehalose. We attribute the differences in preservation to two main factors: difference in the ratio of amount of preservative material to DNA material (0.1M vs 0.02M trehalose), and the different amount of DNA material stored (350 ng vs 24 ng)^15, 16^. We hypothesize the former also explains the difference observed between the DNAStable samples aged at UW and ETH-Z. Nonetheless, the relative rankings of the methods performed at ETH-Z are the same as the rankings found at UW.

It is interesting to note that all preservation methods including trehalose (Trehalose, Sugar Mix) performed well, and exhibit concentration-dependent behavior as illustrated by **Fig. 3b**: the greater the concentration of trehalose, the slower the DNA degrades. Note that DNAStable and GenTegra are comprised of proprietary mixtures and the presence or absence of trehalose is unknown.

### Sequence Analysis of DNA Material

The samples aged at UW were the only samples to be sequenced. All samples were prepared for sequencing prior to aging so that no perturbation of the library (e.g., PCR, ligation steps) were necessary after aging. Each sample had a unique index (i.e., tag) that allowed the identical sequences to be sequenced in the same Illumina NextSeq sequencing run to minimize quality score variation.

First, we explored the difference in observed error rates. We compared the rates of insertions, deletions, and substitutions between all preservation methods at different temperature and time points and between the two files used. As shown in **Fig. 4**, we found minimal variation between the storage methods, with no practical difference between methods and/or files for insertion and deletion errors. Even when looking intently at the substitution rate, which has the most variation, we still observed a maximum difference of just over one percent, which is not practically significant.

**Figure 4.**
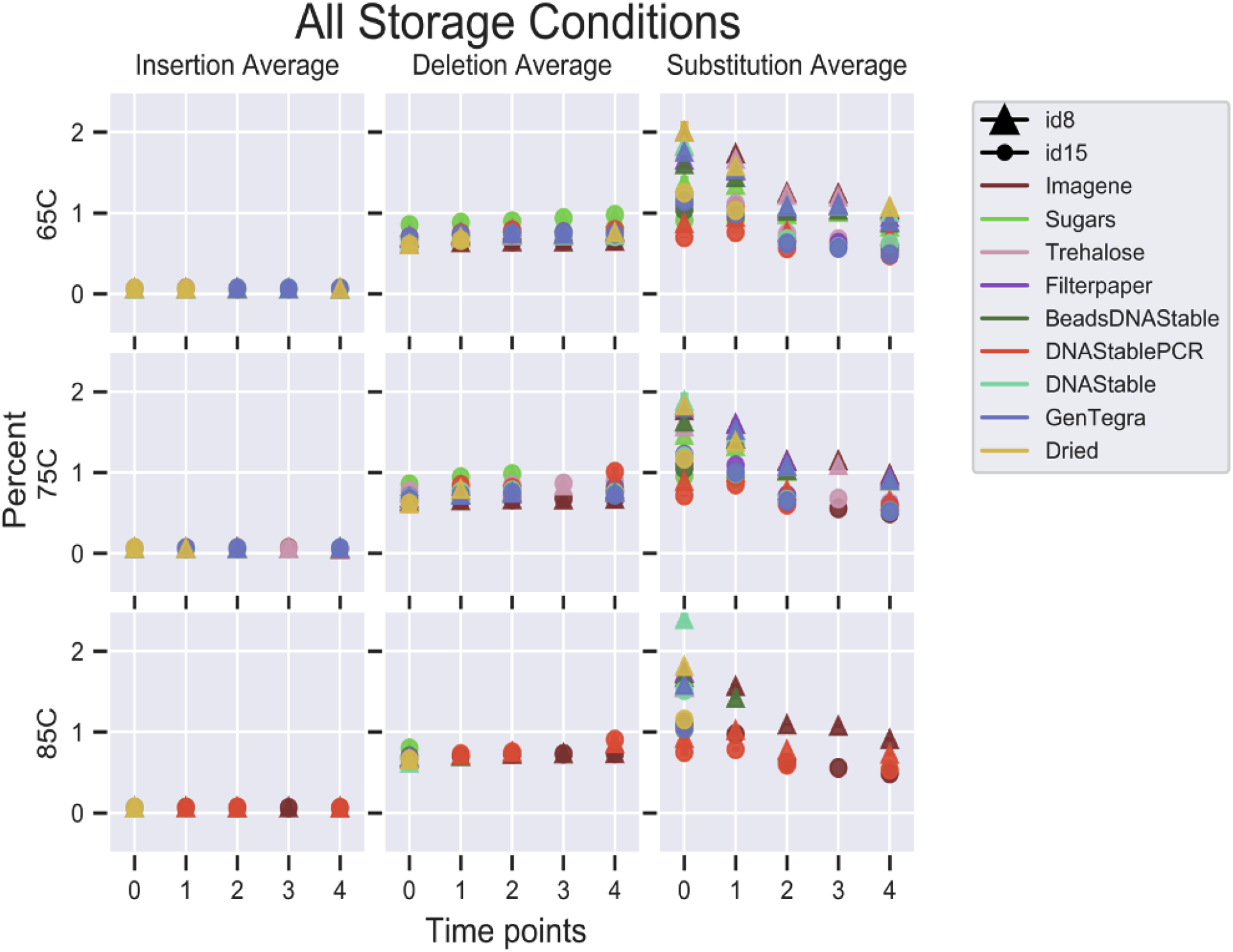
An overview of the error rates observed from sequencing. The error bars are the standard deviation of the method’s triplicates for each time point sequenced. Note that ETH-Z samples were not sequenced. When a certain time point is not plotted, there was either no data for that time point or the file’s average coverage is under the lowest tolerated threshold of 14x (see **Supplemental Section 2** for more information).

More importantly, there is no preservation method that consistently had more or less errors than the others except for the method that employed PCR prior to sequencing (we hypothesize that PCR is, in effect, selecting the most intact and thus less error-prone strands for replication). In fact, as shown at time point 0, there is already a significant amount of variation prior to any aging, and furthermore those initial orderings of methods’ errors do not predict the subsequent method rankings. There was no particular storage method(s) that showed more or fewer errors than other methods across the different temperatures and time points, which suggests that insertion, deletion, and substitution errors are independent from the storage method (see **Supplemental Section 3** for more analysis).

We next explored the relationship between missing sequences in the files between the aged triplicates at different time points to determine if there was sequence-dependent degradation. We examined both the total number of sequences missing from sequencing at each time point as well as which individual sequences were missing. If the total number of sequences missing increased after the pre-aging time point 0, we could hypothesize that there was some sequence-dependent degradation as more-vulnerable sequences degraded. However, we observed no difference in the number of sequences missing across all time points (**Fig. 5a**). This suggests that sequence loss is stochastic across all storage methods.

**Figure 5.**
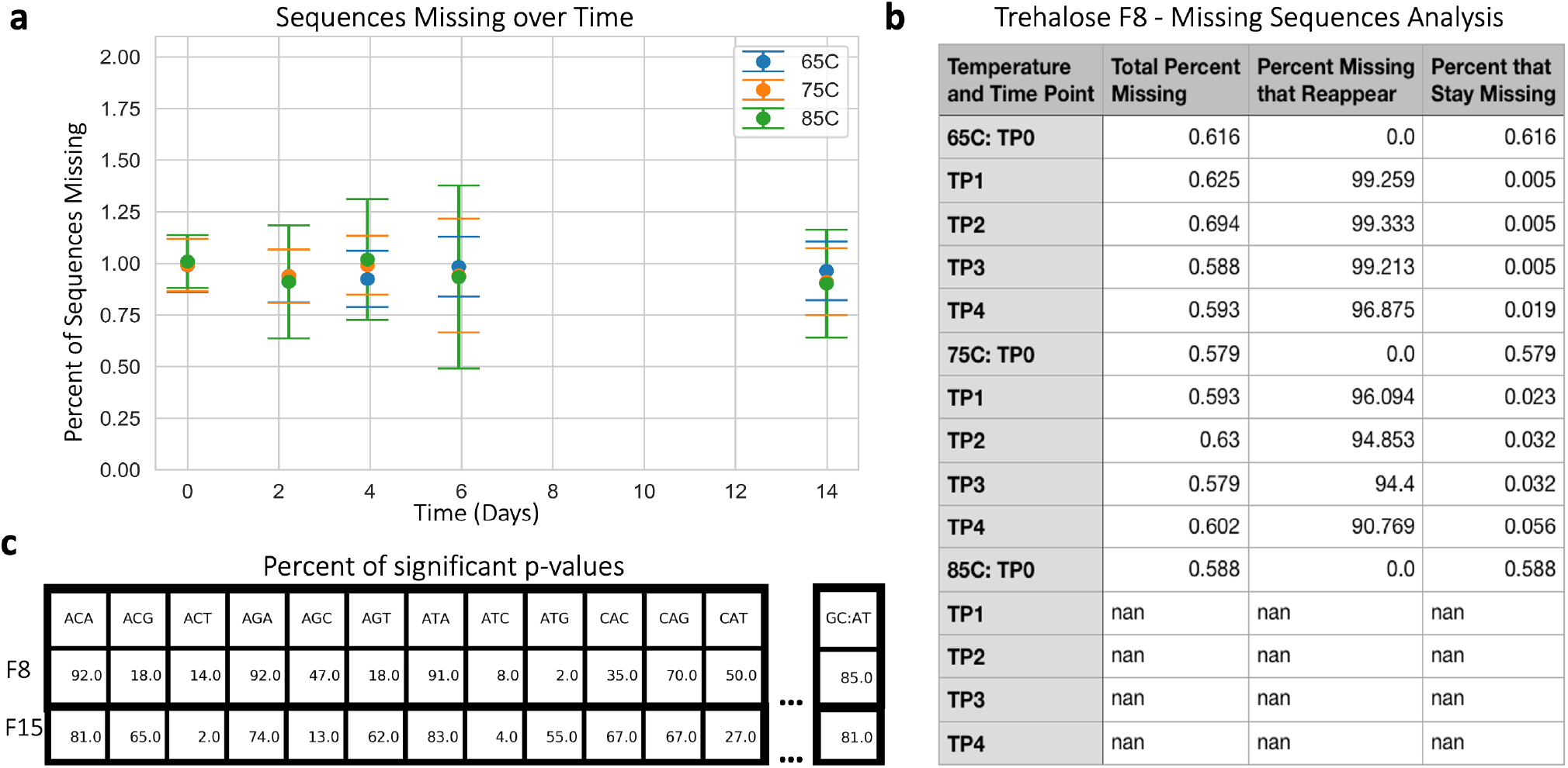
An overview of the sequencing analysis. (a) The mean number of DNA sequences missing from sequencing data over time with error bars representing the standard error of the mean. (b) An example of sequence loss behavior. The “Total Percent Missing” is the mean percentage of the sequences not recovered from each time point and temperature for File 8. Of the sequences missing from each time point and temperature, a non-zero number of sequences typically reappear in subsequent time points, and that percentage is given in “Percent Missing that Reappear”. “Percent that Stay Missing” reports the percentage of total sequences in the file that stay missing. Data for all conditions and files can be found in the Supplemental Files ending with “missing sequences analysis.csv”. (c) A portion of the trimer and GC content analysis giving the percent of samples that had a statistically significant difference in trimer composition between the top 5% most present sequences and the bottom 5% sequences, including missing sequences. For all trimers data for both files, see **Supplemental Section 3**.

This is further supported by our analysis of each individual sequence missing. If a sequence was missing from one time point due to the nature of the sequence being prone to degradation over time, we would not expect to see that sequence again in all subsequent sequencing runs, or in any of the triplicates. However, when we observe a missing sequence, it is often sequenced at a later time point, shown in **Fig. 5b**(as detailed in **Supplemental Section 4.2**). We hypothesize this is due to the stochastic nature of subsampling the sample for sequencing, rather than systematic bias against particular sequences.

That sequence loss is stochastic is yet further supported by results from a trimer and GC content analysis, in which we looked at the prevalence of each trimer (ACA, AGA, ATA, etc.) and the proportion of the sequence that is comprised of guanine (G) and cytosine (C). We found no significant difference between trimer or GC compositions of sequences missing/lowly present and sequences highly present, as shown in **Fig. 5c**. More details can be found in **Supplemental Section 3**.

### Density

Each preservation method in our study improved the half-life of DNA when compared to dehydrated DNA stored with no additives, but at a cost to the physical DNA density of the sample and, at times, simplicity of DNA recovery. Though physical overhead (i.e., molecules added to the sample to preserve DNA) of each technique and time required to extract the DNA may not be significant for many DNA applications, high physical overhead is less appealing for data storage where DNA sequences per volume translates to bytes per volume, and extraction complexity of the DNA limits the rate at which files can be processed. That is to say, the more DNA that can be stored in a given storage container and the faster the samples can be processed, the higher the DNA data density of that container and the higher the throughput.

The highest DNA density was ETH-Z Magnetic NP preservation technique at 3.4 wt% of DNA to encapsulant, though it required the most complex protocol with roughly a 10-20 minute DNA release time ^17^. For methods that involved mixing or simply drying the DNA files with preservatives (Trehalose/ Sugars/ /GenTegra/DNAStable, and Filter Paper), release times are under a minute; however, the DNA loading in the Trehalose and Sugar mix drastically reduced to 0.13 wt% and 0.095 wt%, respectively. Due to the proprietary formulations of GenTegra and DNAStable, we could not accurately calculate the DNA loading but estimate the mix to be similar to the former. DNA stored by mixing or drying the DNA with preservatives degrade significantly faster than the DNA stored in Imagene DNAshells. DNAshells presently have the highest physical overhead due to the borosilicate glass insert and stainless steel shell and cap, though the physical overhead could be reduced as the company develops smaller tubes to decrease physical overhead.

## Discussion

DNA samples aged in preservation methods without exposure to any water or air (Imagene) were found to decay much slower than samples exposed to water and air, which supports previous findings^7, 10, 14,18^. Samples preserved with trehalose exhibited concentration dependence, with the higher concentration of trehalose conferring more protection against degradation (**Fig. 3b**, and the amount of DNA stored per sample and the ratio of DNA to storage molecules may also play an important role on the rate of degradation, depending on the preservation method used^15, 16^.

Depending on the storage method, there are sometimes significant changes in the DNA encapsulation and retrieval process. Imagene DNAShells (Imagene) and magnetic nanoparticles (ETH-Z Magnetic NP) were the most complex storage methods to retrieve DNA from, as they either required specialized equipment or involved several steps. In contrast, DNA combined with preservatives and simply dehydrated have the simplest recovery procedures for they only require a simple re-hydration protocol. This leads to an inverse relationship between stability and process complexity. Surprisingly, DNA stored in the presence of trehalose or GenTegra did not fit this storage complexity trend and instead degraded much slower than anticipated.

There was a real concern that half-life values would not be the only difference between storage methods, and that the sequences would accumulate insertion, deletion, or substitution errors at varying rates. However, we observed that error rates did not differ with any practical significance between the preservation methods. Furthermore, within the tight constraints of the sequences examined here (i.e., no homopolymers in the payload region, balanced GC content), sequence degradation was found to be stochastic. This is encouraging to the field of DNA data storage, where stochastic errors and sequence erasures are already dealt with easily with various means of error correction such as Reed-Solomon codes^4, 7, 19^.

Automation will very likely be the key in obtaining a scalable DNA data storage system. It is foreseeable that a solution containing DNA files and a preservative could be mixed together and dried on a thin film for storage. When the file needs to be accessed, it would be re-hydrated for a short time (likely on the order of a few seconds) and then processed for sequencing. Several early examples of this already exist on a glass top plate using a digital microfluidic system^16, 20^ but could extend to other liquid dispensing systems such as acoustic or ink-jet dispensing. Development of an automated DNA storage and retrieval system would require thorough investigation of the effects of sampling and preserving the same sample multiple times, as studies in genomic DNA have shown this causes a non-negligible effect on recovery and may be preservation material dependent^21^.

For a more comprehensive guide to DNA degradation, more long-term studies should be conducted in which DNA is allowed to degrade for years at a time, rather than days or a few weeks as most aging experiments have done, and at concentrations and temperatures more reflective of storage conditions users anticipate. This is because DNA decay depends on storage material, temperature, and other factors^11, 22^.

Yet it is important to note that two different labs using different experimental aging setups and different length sequences produced the same relative ranking of those methods, though the half-life values did differ. This underscores the fact that exact half-life values are difficult to extrapolate and very sensitive to the conditions used to achieve them. We encourage a more robust examination of all these variables depending on a user’s end goals and likely storage conditions.

Users of future DNA data storage systems will require a broad range of stability, from hundreds to thousands of years, but the error analysis results presented here demonstrate the interchangeable nature of all the methods examined regarding sequencing data quality. As this work shows, users must choose their preferred storage method based on their desired half-life, data density, and process complexity.

## Methods

### DNA Material Stored

It is important to note there are slight differences between the storage protocols used by UW and ETH. The protocol used by UW is shown in **Fig. 1**. 28,974 unique sequences of dsDNA 150nt in length were prepared for NextGen sequencing for a final length of 310 base pairs. Storage experiments performed by ETH contained only 7,373 of the DNA sequences (File 15) and did not prepare them for NextGen sequencing, so the length of DNA stored in their ovens was 150nt.

### Sample preparation

From 150mer DNA pools synthesized by Twist Biosciences, two files were amplified. File 8 was comprised of 21,601 unique DNA sequences while File 15 was comprised of 7,373 unique DNA sequences. Within each file, the primer sequences (first and last twenty bases of the 150mer) are conserved and the middle 110 bases are distributed such that there are no homopolymers and the GC content is approximately 50%.

Each of the files was PCR amplified in multiple rounds to minimize PCR bias (when there is an uneven distribution of each DNA sequence due to the stochastic nature of PCR) using the following recipe: 5 *μ*L of 1 ng/*μ*L of DNA template, 2.5 *μ*L of each primer at 10*μ*M, 50 *μ*L of Kappa HiFi 2x, and 40 *μ*L of molecular grade water. The PCR protocol was then: (1) 95C for 3 min, (2) 98C for 20 sec, (3) 62C for 20 sec, (4) 72C for 15 sec, (5) repeat steps 2-4 a total of 13 or 20 times for files 8 and 15 respectively,(6) 72C for 30 sec.

The samples were then PCR amplified following the same protocol, but with forward primes now having a 25N overhang to allow for sequencing (this is an artifact of the Illumina NextSeq, as largely uniform sequences at the beginning of sequencing do not allow for accurate cluster calling and so the first 25 bases being completely random solves this problem).

The samples were then quantified with qPCR, and then pooled together so that File 8 had approximately twice as many copies of each sequence as File 15.

This pool of File 8 and File 15 sequences was then split into 96 identical alequots and prepared for aging and sequencing with ligation. (Note that in actuality-9 of these alequots that were to become the basis for DNAStable + PCR also had 323,875 extra sequences added to its sample to make the PCR conditions slightly more complex, as would be more realistic in a typical setting, though the number of copies of each sequence remained the same as the other samples.) Ligation was done with a modified version of Illumina TruSeq Nano ligation protocol and TruSeq ChIP Sample Preparation protocol. Step by step instructions are in **Supplementary Section 5** for convenience, but briefly, samples were first converted to blunt ends with the ERP2 reagent and directions provided in the Illumina TruSeq Nano kit, then purified with AMPure XP beads according to the TruSeq ChIP protocol. An ‘A’ nucleotide was added to the 3‘ ends of the blunt DNA fragments with the TruSeq Nano’s A-tailing ligase and protocol, followed by ligation to the Illumina sequencing adapters with the TruSeq Nano reagents and protocol. We then cleaned the samples with Illumina sample purification beads and enriched the sample using an 8-cycle PCR protocol given in the TruSeq Nano protocol.

For the enrichment, all samples (each now with a unique ligation index) were enriched using the following recipe: 3*μ*L of a ligation sample, 3*μ*L of the PCR Primer Cocktail provided in the TruSeq Nano kit, 12*μ*L of Enhanced PCR Mix provided in the TruSeq Nano kit, and 12*μ*L of molecular grade water. The following PCR protocol was used: (1) 95C for 3 min, (2) 98C for 20 sec, (3) 60C for 15 sec, (4) 72C for 30 sec, (5) repeat steps 2–4 for a total of eight times. The length of enriched products was confirmed using a Qiaxcel bioanalyzer. Reformatted instructions are given in **Supplementary Section 5** for convenience.

After enrichment, the PCR products with the same ligation index were pooled together and PCR purified with a QIAquick PCR purification kit with molecular grade water at the elution step.

Each sample was then quantified with a Qubit 2.0 fluorometer and each preservation method took 350 ng of material per sample. The details of each preservation method are below.

### No Additives

From samples described in the “Sample preparation” section, nine samples, each with a unique ligation index, were used for this preservation method. From each ligation index, five alequots of 350ng were made and placed in 0.6mL eppindorf tubes (one alequot for each time point). Each tube was then dehydrated in a SPD 1030 speedvac on high.

After aging, nine samples (one alequot from each index, in other words, three indices per temperature) were removed from their respective heat sinks and allowed to come to room temperature, then rehydrated with 50*μ*L of molecular grade water, capped, and vortexed on a benchtop vortexer. The samples then sat at room temperature for 20 minutes prior to quantification with qPCR (see Methods section, “Quantifying Degradation with qPCR”).

### DNAStable

From samples described in the “Sample preparation” section, nine samples, each with a unique ligation index, were used for this preservation method. From each ligation index, five alequots of 350ng, with a volume ranging from 3.4-5*μ*L, were made and placed in 0.6mL eppindorf tubes (one alequot for each time point). Each tube then had 20*μ*L of DNAStable LD (product number 53001-066) added, and the mixture was pipetted up and down three times. Each tube was then dehydrated in a SPD 1030 speedvac on high.

After aging, nine samples (one alequot from each index, in other words, three indices per temperature) were removed from their respective heat sinks and allowed to come to room temperature, then rehydrated with 50*μ*L of molecular grade water, capped, and vortexed on a benchtop vortexer. The samples then sat at room temperature for 20 minutes prior to quantification with qPCR (see Methods section, “Quantifying Degradation with qPCR”).

### DNAStable PCR

The preservation and rehydration protocol is identical to DNAStable.

For DNAStable PCR, the samples were analyzed with qPCR in an identical manner to the other methods, then File 8 and File 15 were accessed with PCR (see **Supplemental Section 1.1** for primer sequences), re-ligated with the same protocol described above (see **Supplemental Section 1.2**), and sequenced using the same protocol as all other methods. The amplification protocol for File 8 and File 15 was as follows:

First, amplify each sample using the primers listed in **Supplemental Section 1.1**. For time point 0, 0.2*μ*L of sample was used and 10 cycles were performed. For time point 1, 8*μ*L of sample was used and 12 cycles were performed for the 65C samples, while 16 cycles were performed for the 75C and 85C samples. In addition to the sample DNA, there were 10*μ*L of 2x Kapa HiFi, 1*μ*L of each primer at 10*μ*M, and 7*μ*L of molecular grade water. The thermocycle protocol was: (1) 95C for 3 min, (2) 98C for 20 sec, (3) 62C for 20 sec, (4)72C for 15 sec, (5) repeat steps2-4 for a total of the number of times stated at the beginning of this paragraph.

Then, amplify the resulting sample again except that now there is a 25Nmer (25 random nucleotides) on the 5’ end of the forward primer. Follow the same protocol as above, but now there are always 12 cycles and 1*μ*L of DNA product.

Then, rather than performing ligation in preparation for sequencing, perform one last PCR with appropriate overhanging sequences. Follow the same PCR protocol as the previous paragraph, but this time using primers containing each files’ primer sequence, as well as the relevant index sequence and Illumina adapter and sequencing primer, as listed in **Supplemental Section 1**.

### Mag-Bind DNAStable

From samples described in the “sample preparation” section, nine samples, each with a unique ligation index, were used for this preservation method. From each ligation index, five aliquots of 350 ng, with volumes ranging 20-30*μ*L, were made and placed in 0.6mL eppendorf tubes (one aliquot for each time point). Each tube than had 1.2X ratio of Mag-Bind beads to DNA volume added, and the mixture was vortexed and allowed to sit at room temperature for 5 minutes. After magnetic bead separation, the supernatant was discarded. Then the samples were washed with 200 uL of 70% ethanol, allowed to sit for 1 minute before separating the beads and discarding the supernatant (performed 2x). Any residual ethanol was air dried for 10 minutes. Each tube then had 20*μ*L of DNAStable LD (product number 53001-066) added, and the mixture was vortexed to mix. Each tube was then dehydrated in a SPD 1030 speedvac on high.

After aging, nine samples (one alequot from each index, in other words, three indices per temperature) were removed from their respective heat sinks and allowed to come to room temperature, then rehydrated with 50*μ*L of molecular grade water, capped, and vortexed on a benchtop vortexer. The samples then sat at room temperature for 20 minutes before magnetic bead separation. The supernatant was then used for qPCR quantification (see Methods section, “Quantifying Degradation with qPCR”).

### Sugar Mix

From samples described in the “sample preparation” section, nine samples, each with a unique ligation index, were used for this preservation method. From each ligation index, five aliquots of 350 ng, with volumes ranging 7-9*μ*L, were made and placed in 0.6mL eppendorf tubes (one aliquot for each time point). Each tube then had equal volume*μ*L of 0.2M Sugar Mix (0.1 M trehalose, 0.05 M raffinose, 0.05 M mannitol, 0.125 mM uric acid) added and the mixture was pipetted up and down three times. Each tube was then dehydrated in a SPD 1030 speedvac on high.

After aging, nine samples (one alequot from each index, in other words, three indices per temperature) were removed from their respective heat sinks and allowed to come to room temperature, then rehydrated with 50*μ*L of molecular grade water, capped, and vortexed on a benchtop vortexer. The samples then sat at room temperature for 20 minutes prior to quantification with qPCR (see Methods section, “Quantifying Degradation with qPCR”).

### Trehalose

From samples described in the “sample preparation” section, nine samples, each with a unique ligation index, were used for this preservation method. From each ligation index, five aliquots of 350 ng, with volumes ranging 7-9*μ*L, were made and placed in 0.6mL eppendorf tubes (one aliquot for each time point). Each tube then had equal volume*μ*L of 0.2M Trehalose added, and the mixture was pipetted up and down three times. Each tube was then dehydrated in a SPD 1030 speedvac on high.

After aging, nine samples (one alequot from each index, in other words, three indices per temperature) were removed from their respective heat sinks and allowed to come to room temperature, then rehydrated with 50*μ*L of molecular grade water, capped, and vortexed on a benchtop vortexer. The samples then sat at room temperature for 20 minutes prior to quantification with qPCR (see Methods section, “Quantifying Degradation with qPCR”).

### GenTegra

From samples described in the “Sample preparation” section, nine samples, each with a unique ligation index, were used for this preservation method. From each ligation index, five alequots of 350ng, with volumes ranging 6-15*μ*L, were made and placed in GenTegra tubes (one alequot for each time point). This mixture was then pipetted up and down 10 times as per GenTegra’s instruction. The GenTegra tubes were 0.5mL with GenTegra’s proprietary mixture at the bottom of each tube (product number GTD2100-S). Each tube was then dehydrated in a SPD 1030 speedvac on high.

After aging, nine samples (one alequot from each index, in other words, three indices per temperature) were removed from their respective heat sinks and allowed to come to room temperature, then rehydrated with 50*μ*L of molecular grade water, capped, and vortexed on a benchtop vortexer. The samples then sat at room temperature for 20 minutes prior to quantification with qPCR (see Methods section, “Quantifying Degradation with qPCR”).

### Filter Paper

From samples described in the “Sample preparation” section, nine samples, each with a unique ligation index, were used for this preservation method. From each ligation index, five alequots of 350ng, with volumes ranging 6-10*μ*L, were pipetted 2*μ*L at a time onto 2.5mm diameter VWR Grade 415 filter paper (product number 28320-121) and dried at 30C between rounds of pipetting. The final, dry 2.5 diameter circles of filter paper were then individually placed in 0.6mL eppindorf tubes.

After aging, nine samples (one alequot from each index, in other words, three indices per temperature) were removed from their respective heat sinks and allowed to come to room temperature, then rehydrated with 50*μ*L of molecular grade water, capped, and vortexed on a benchtop vortexer with the filter paper still in the tube. The samples then sat at room temperature for 20 minutes prior to quantification with qPCR (see Methods section, “Quantifying Degradation with qPCR”).

### Imagene

From samples described in the “Sample preparation” section, nine samples, each with a unique ligation index, were used for this preservation method. Each sample was then shipped overnight on dry ice to Imagene with two extra samples prepared the same as the others, but only used as shipping controls. Imagene then deposited 350 ng of the shipped DNA material into each DNAshell. The samples were then shipped back to the UW with the shipping controls. The samples were stored at −20C until the aging process started.

After aging, nine samples (one alequot from each index, in other words, three indices per temperature) were removed from their respective heat sinks and allowed to come to room temperature. The metal DNAshells were then pierced with the included shell piercer, and then rehydrated with 50*μ*L of molecular grade water and pipetted up and down 10 times. The samples then sat at room temperature for 20 minutes and were transferred out of DNAshells to PCR tubes prior to quantification with qPCR (see Methods section, “Quantifying Degradation with qPCR”).

### No Additives - Performed by ETH-Z

A sample of F15 and primers were shipped to ETH-Z. F15 was PCR amplified and purified using QIAquick PCR purification Kit (Qiagen). Following elution with MiliQ water, the final concentration of DNA was 12 ng/uL.

Five aliquots of 24 ng (2*μ*L), were made and placed in 2 mL eppendorf tubes (one aliquot for each time point). Each tube was then dehydrated in a vacuum centrifuge for 2 hours at 60C and aged in a desiccator containing a saturated NaBr solution within an oven. After aging, samples were removed from the oven and allowed to come to room temperature. They were then rehydrated and quantified with qPCR.

### Trehalose - Performed by ETH-Z

Five aliquots of 24 ng (2*μ*L), were made and placed in 2 mL eppendorf tubes (one aliquot for each time point). Each tube then had 4 uL of 0.02M Trehalose solution added and was mixed. Each tube was then dehydrated in a vacuum centrifuge for 2 hours at 60C and aged in a desiccator containing a saturated NaBr solution within an oven. After aging, samples were removed from the oven and allowed to come to room temperature. They were then rehydrated and quantified with qPCR.

### DNAStable - Performed by ETH-Z

Five aliquots of 24 ng (2*μ*L), were made and placed in 1.5 mL eppendorf tubes pre-coated with DNAStable (one aliquot for each time point). Each tube was then dehydrated in a vacuum centrifuge for 2 hours at 60C and aged in a desiccator containing a saturated NaBr solution within an oven. After aging, samples were removed from the oven and allowed to come to room temperature. They were then rehydrated and quantified with qPCR.

### Magnetic Nanoparticles - Performed by ETH-Z

Samples were encapsulated and deencapsulated in magnetic nanoparticles following the protocol reported in Chen et al.

Five aliquots of 40*μ*L final particle solution were made and placed in 1.5 mL eppendorf tubes (one aliquot for each time point). Each tube was then dehydrated in a vacuum centrifuge overnight at 30C and aged in a desiccator containing a saturated NaBr solution within an oven. After aging, samples were removed from the oven and allowed to come to room temperature.

### Aging DNA

After preservation, all samples except time point 0 samples were placed in their respective ovens concurrently and kept uncapped at 50% relative humidity (except for the sealed Imagene DNAshell capsules).Relative humidity was controlled through a well-established technique using saturated salt solutions^23^. Sodium bromide was dissolved in water until it was no longer dissolvable with percipitate to create a saturated solution of sodium bromide. A deep 1L petri dish was filled with the saturated salt solution and placed on the bottom oven rack to maintain 50% RH atmosphere at the three different temperatures.

The ovens used were Quincy Lab, Inc. Model 10E-LT Lab Oven, with form-fitted insulation over the body of the oven above the control system made of polyisocyanurate rigid foam insulation board that was one inch thick to prevent condensation.

An aluminum heat sink (a one inch thick block of aluminum) with holes the width and depth of each tube was milled and placed in each of the three ovens.

In each oven, a thermometer was placed through the vent and rested on the heat sink and monitored daily for temperature fluctuations greater than 2 degrees, which were not observed during the experiment except immediately after removing the heat sinks at each time point.

### Quantifying Degradation with qPCR

After each aging time point, samples were diluted 1:100 with molecular water and the amount of full-length product present was quantified with qPCR. The standard used was an ultramer ordered from IDT, whose sequence is given in **Supplemental Section 1**. The qPCR recipe is as follows: 1*μ*L DNA sample, 0.5*μ*L of each post-ligation primer (given in **Supplemental Section 1**) at 10*μ*M, 10*μ*L Kapa HiFi, 7*μ*L molecular water, 1*μ*L 20x Eva Green. The qPCR thermocycling protocol was: (1) 95C for 30 sec, (2) 98C for 20 sec, (3) 60C for 20 sec, (4) 72C for 30 sec, (5) repeat steps 2-4 39 times. A negative control was included on each plate that had no DNA sample included with the mixture of all other reagents.

## Data Availability Statement

The data from this study are available from the corresponding authors upon reasonable request.

## Code Availability Statement

Code used to do the analysis presented in the paper is available from the corresponding authors upon reasonable request. Encoding and decoding code is proprietary.

## Acknowledgements

Work performed at UW was jointly funded by Microsoft and the DARPA Molecular Informatics Program. Work performed at ETH-Z was funded by Microsoft.

## Author contributions statement

L.O. designed, performed, and analyzed experiment data and wrote the manuscript. B.H.N. designed and analyzed experiment data and wrote the manuscript. R.M. analyzed sequencing data. W.D.C. and A.X.K. performed and analyzed experiments. S.D.A. designed sequencing error analysis methods and performed sequencing analysis. R.N.G. analyzed and discussed data. L.C. and K.S. directed and supervised the work.

