## Supplemental Information for "A empirical comparison of preservation methods for synthetic DNA data storage" (from another version of this preprint: version 1, 2020-09-19)

### A comprehensive study of synthetic DNA preservation for DNA data storage

Supplementary Material

Organick *et al.*

### Contents

**1 Performing and Analyzing qPCR** **3**
- 1.1 Primer Sequences . . . . . . . . . . 3
- 1.2 Primer Sequences for the DNAStable PCR Samples . . . . . . . 3

**2 Determining a Coverage Threshold** **6**
- 2.1 Insertion/Substitution/Deletion Rates as a Function of Coverage 6

**3 Sequence behavior analysis** **9**

**4 Overview Table** **18**

**5 Ligation Protocol** **19**

**6 Arrhenius Equation** **23**

### 1 Performing and Analyzing qPCR

#### 1.1 Primer Sequences

Each file has a unique primer pair, facilitating random access in DNA data storage. The primer sequences are in **Supplementary Table 1**.

| File | Forward Primer | Reverse Primer |
|---|---|---|
| ID8 | TTCGTTCGTCGTTGATTGGT | ACAAACTCATGGCTCCGTTT |
| ID15 | ACATTCCGTGCCATTGGATT | TTTGTGGAACGATTTGCCGA |
| Post-Ligation | AATGATACGGCGACCACCGA | CAAGCAGAAGACGGCATACG |

Supplementary Table 1: The forward and reverse primers needed to amplify each file, as well as the primer pair used to enrich the sequences after ligation, but prior to aging.

The ultramer used to quantify the UW (non-ETH) samples after degradation utilized the post-ligation primers. The ultramer sequence is:

5' AATGATACGGCGACCACCGAGATCTTGACCAAACACTCTTTCCC-TACACGACGCTCTTCCGATCTACTGA CTGAGTACGACATGCATGCAGC-CGTCCATAGCCTTGTTCGTTGTCATAGTATGTAGCTACTGCACGTAT GCAT-ACTGAGTCTGTACATGAGTAGTGCACAGTAGTAGACGCGTCATCAGAG-TATGCATATCAGCATCG CGTAGCTAGCTAGGCGGAAACGTAGTGAAG-GTAGATCGGAAGAGCACACGTCTGAACTCCAGTCACTGA CCAATCTCG-TATGCCGTCTTCTGCTTG 3'

#### 1.2 Primer Sequences for the DNAStable PCR Samples

Here, **Supplementary Tables 2** and **3** show the overhanging PCR primers used to prepare samples in the *DNAStable PCR* aging condition after random access, prior to sequencing. Thus, the samples did not have to be ligated again.

The spaces within the sequences listed in the tables are to help the reader separate the different domains of the primer sequence, which are: Illumina universal flow cell adapter, Illumina index, Illumina sequencing primer, file 8/15 primer.

| Primer Description | Sequence |
|---|---|
| Forward, ID8, index H | AATGATACGGCGACCACCGAGATCTACACGTACTGACA CACTCTTTCCCTACACGACGCTCTTCCGATCT TTCGTTCGTCGTTGATTGGT |
| Reverse, ID8, index H4 | CAAGCAGAAGACGGCATACGAG GGAATCTC GTGACTG-GAGTTCAGACGTGTGCTCTTCCGATC ACAAACTCATG-GCTCCGTTT |
| Reverse, ID8, index H5 | CAAGCAGAAGACGGCATACGAG TTCTGAAT GTGACTG-GAGTTCAGACGTGTGCTCTTCCGATC ACAAACTCATG-GCTCCGTTT |
| Reverse, ID8, index H6 | CAAGCAGAAGACGGCATACGAG ACGAATTC GTGACTG-GAGTTCAGACGTGTGCTCTTCCGATC ACAAACTCATG-GCTCCGTTT |
| Reverse, ID8, index H7 | CAAGCAGAAGACGGCATACGAG AGCTTCAG GTGACTG-GAGTTCAGACGTGTGCTCTTCCGATC ACAAACTCATG-GCTCCGTTT |
| Reverse, ID8, index H8 | CAAGCAGAAGACGGCATACGAG GCGCATTA GTGACTG-GAGTTCAGACGTGTGCTCTTCCGATC ACAAACTCATG-GCTCCGTTT |
| Reverse, ID8, index H9 | CAAGCAGAAGACGGCATACGAG CATAGCCG GTGACTG-GAGTTCAGACGTGTGCTCTTCCGATC ACAAACTCATG-GCTCCGTTT |
| Reverse, ID8, index H10 | CAAGCAGAAGACGGCATACGAG TTCGCGGA GTGACTG-GAGTTCAGACGTGTGCTCTTCCGATC ACAAACTCATG-GCTCCGTTT |
| Reverse, ID8, index H11 | CAAGCAGAAGACGGCATACGAG GCGCGAGA GTGACTG-GAGTTCAGACGTGTGCTCTTCCGATC ACAAACTCATG-GCTCCGTTT |
| Reverse, ID8, index H12 | CAAGCAGAAGACGGCATACGAG CTATCGCT GTGACTG-GAGTTCAGACGTGTGCTCTTCCGATC ACAAACTCATG-GCTCCGTTT |

Supplementary Table 2: The forward and reverse primers needed to attach the Illumina indices, sequencing primer, and adapters to file 8 via PCR, only utilized for the DNAStable + PCR aging method since these samples were aged, then PCR'ed with the short primers, then prepped for sequencing using these overhang PCR sequences.

| Primer Description | Sequence |
|---|---|
| Forward, ID15, index H | AATGATACGGCGACCACCGAGATCTACACGTACTGACA CACTCTTTCCCTACACGACGCTCTTCCGATCT ACATTC-CGTGCCATTGGATT |
| Reverse, ID15, index H4 | CAAGCAGAAGACGGCATACGAG GGAATCTC GTGACTG-GAGTTCAGACGTGTGCTCTTCCGATC TTTGTGGAAC-GATTTGCCGA |
| Reverse, ID15, index H5 | CAAGCAGAAGACGGCATACGAG TTCTGAAT GTGACTG-GAGTTCAGACGTGTGCTCTTCCGATC TTTGTGGAAC-GATTTGCCGA |
| Reverse, ID15, index H6 | CAAGCAGAAGACGGCATACGAG ACGAATTC GTGACTG-GAGTTCAGACGTGTGCTCTTCCGATC TTTGTGGAAC-GATTTGCCGA |
| Reverse, ID15, index H7 | CAAGCAGAAGACGGCATACGAG AGCTTCAG GTGACTG-GAGTTCAGACGTGTGCTCTTCCGATC TTTGTGGAAC-GATTTGCCGA |
| Reverse, ID15, index H8 | CAAGCAGAAGACGGCATACGAG GCGCATTA GTGACTG-GAGTTCAGACGTGTGCTCTTCCGATC TTTGTGGAAC-GATTTGCCGA |
| Reverse, ID15, index H9 | CAAGCAGAAGACGGCATACGAG CATAGCCG GTGACTG-GAGTTCAGACGTGTGCTCTTCCGATC TTTGTGGAAC-GATTTGCCGA |
| Reverse, ID15, index H10 | CAAGCAGAAGACGGCATACGAG TTCGCGGA GTGACTG-GAGTTCAGACGTGTGCTCTTCCGATC TTTGTGGAAC-GATTTGCCGA |
| Reverse, ID15, index H11 | CAAGCAGAAGACGGCATACGAG GCGCGAGA GTGACTG-GAGTTCAGACGTGTGCTCTTCCGATC TTTGTGGAAC-GATTTGCCGA |
| Reverse, ID15, index H12 | CAAGCAGAAGACGGCATACGAG CTATCGCT GTGACTG-GAGTTCAGACGTGTGCTCTTCCGATC TTTGTGGAAC-GATTTGCCGA |

Supplementary Table 3: The forward and reverse primers needed to attach the Illumina indices, sequencing primer, and adapters to file 15 via PCR, only utilized for the DNAStable + PCR aging method since these samples were aged, then PCR'ed with the short primers, then prepped for sequencing using these overhang PCR sequences.

#### 2 Determining a Coverage Threshold

The purpose of determining a coverage threshold is to exclude files that have so few average reads that the data could inaccurately reflect the number of strands that have been degraded. In other words, a strand that is missing is more likely to be due to random chance of not being read rather than actually having been degraded. To determine a threshold of when to assume that files might more likely be "missing" from random chance rather than being degraded, a plot of average file coverage versus percent missing was created. Only time point zero files (from all storage conditions) were included to determine the coverage threshold. The coverage threshold is the same for both id8 and id15 files. **Supplementary Figure 1** displays how an average coverage of 14 and beyond had a relatively low percent of strands missing from reads:

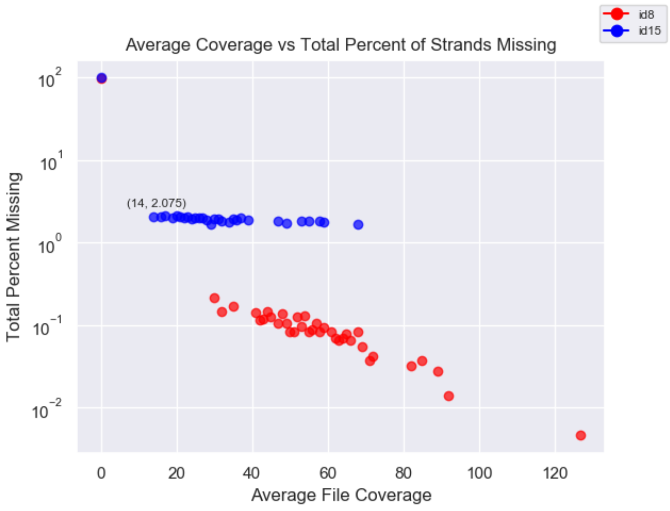

Supplementary Figure 1: Relationship between sequencing coverage and sequences missing.

All files were subsampled to the determined coverage threshold, which is an average of 14 reads per sequence. Every file with greater than or equal to an average coverage of 14 was subsampled 100 times, and the average coverage for each sequence was calculated and used in all subsequent analysis.

##### 2.1 Insertion/Substitution/Deletion Rates as a Function of Coverage

Average coverage versus insertion/deletion/substitution rate were plotted to ensure no correlation between average coverage and error rate. Like with determining coverage threshold, only time point zero data are plotted. We find no practi-

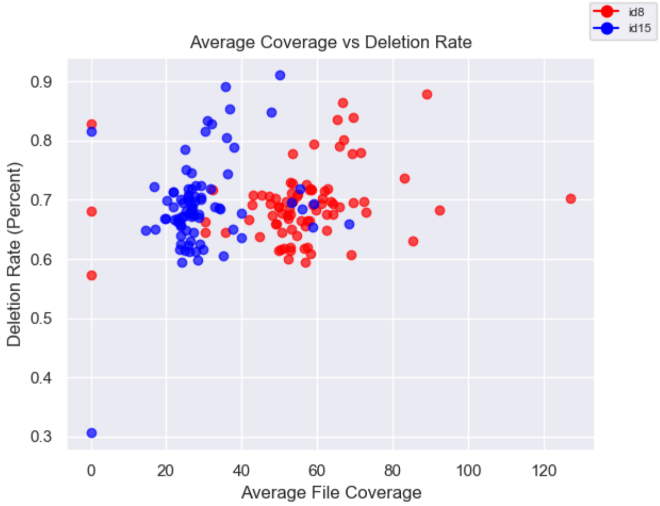

Supplementary Figure 2: Relationship between sequencing coverage and deletion error rate.

cally significant relationship between coverage and error rate (**Supplementary Figures 2, 3, 4**).

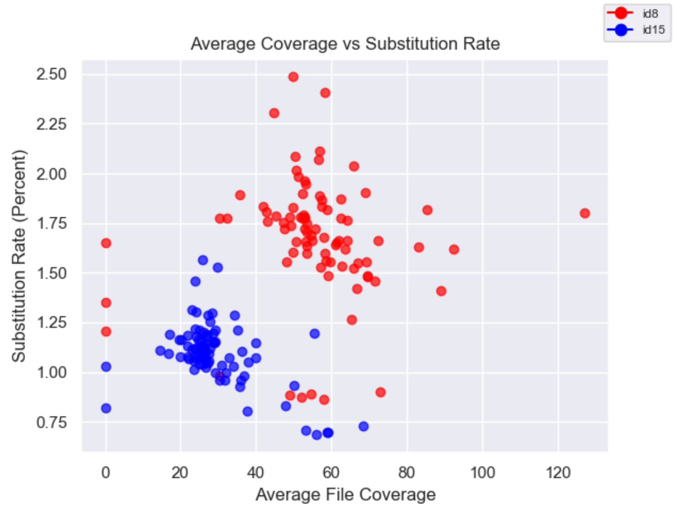

Supplementary Figure 3: Relationship between sequencing coverage and substitution error rate.

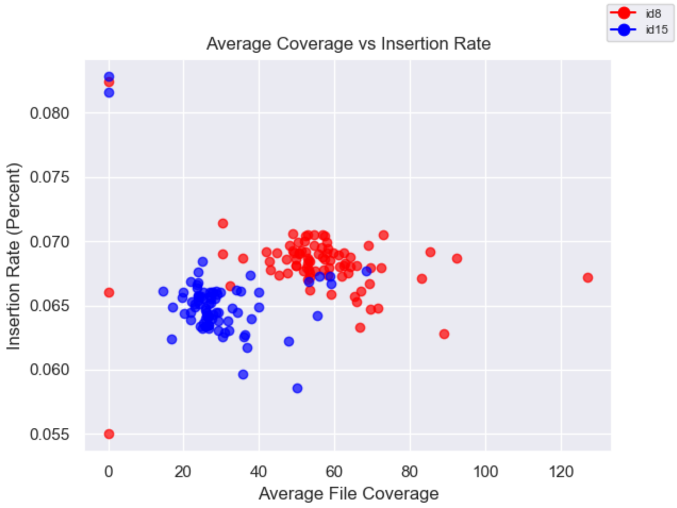

Supplementary Figure 4: Relationship between sequencing coverage and insertion error rate.

### 3 Sequence behavior analysis

Here, we present the sequence behavior observed after next-generation sequencing, as it is important to know if any particular types of sequences degrade faster than others, or if sequence degradation is stochastic in nature.

We analyze both the absence of sequences in varying levels of detail, as well as individual sequence composition in the form of a trimer analysis.

#### 3.1 Percent of sequences missing over time

Here, we show that the percent of sequences missing is not dependent on degradation. All samples that had enough material (as measured by the qPCR) to directly sequence were sequenced.

As shown in the main text, the percent of sequences missing is constant. Here, we present each preservation method individually in Supplementary Figures 5 to 13. Note that several times the data are so similar not all conditions are visible because they are stacked so closely on top of one another.

That the percent of sequences missing constant shows degradation likely does not impact certain sequences more than others, as those hypothetical "degradation-prone" sequences would have degraded faster than the surrounding material and would not have been sequenced.

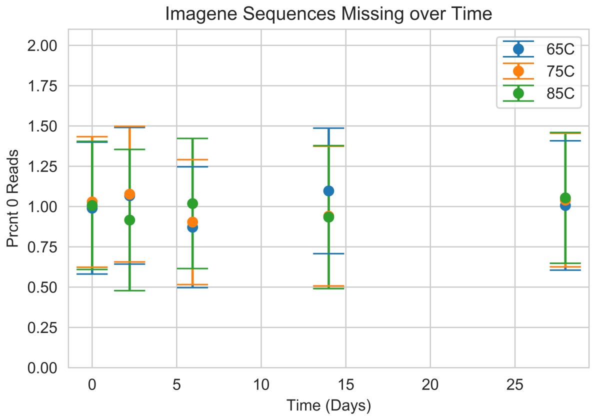

Supplementary Figure 5: Sequences missing from each time point and temperature for all triplicates, where error bars are the standard error from the mean.

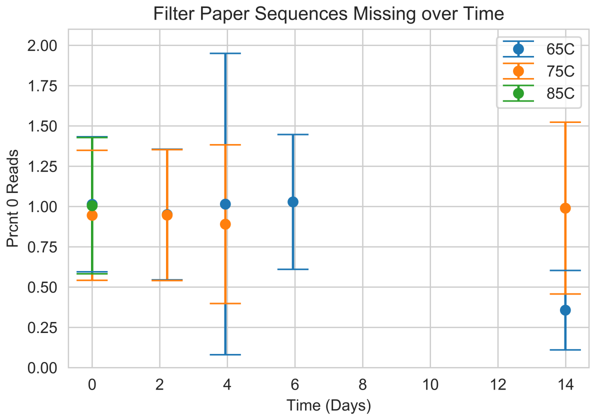

Supplementary Figure 6: Sequences missing from each time point and temperature for all triplicates, where error bars are the standard error from the mean.

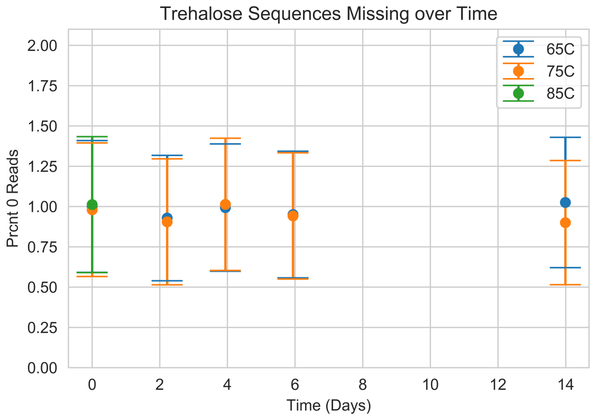

Supplementary Figure 7: Sequences missing from each time point and temperature for all triplicates, where error bars are the standard error from the mean.

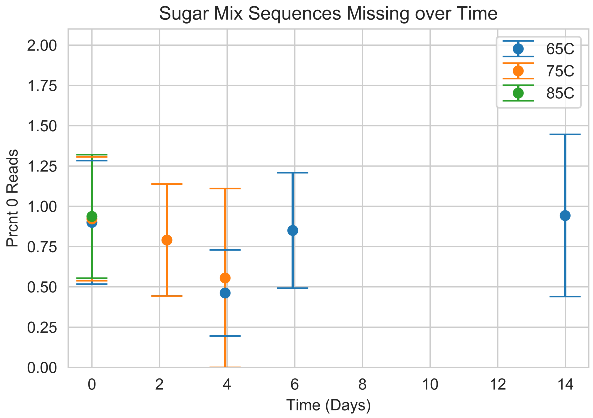

Supplementary Figure 8: Sequences missing from each time point and temperature for all triplicates, where error bars are the standard error from the mean.

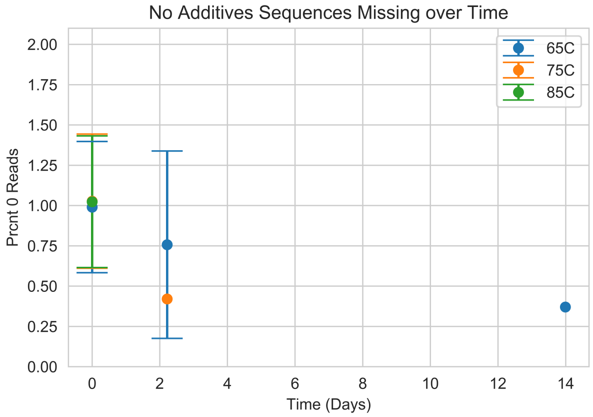

Supplementary Figure 9: Sequences missing from each time point and temperature for all triplicates, where error bars are the standard error from the mean.

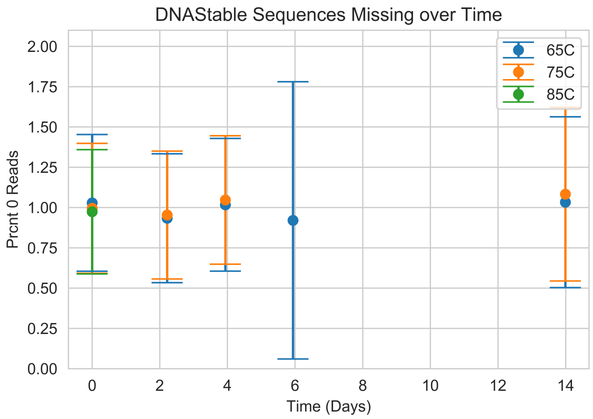

Supplementary Figure 10: Sequences missing from each time point and temperature for all triplicates, where error bars are the standard error from the mean.

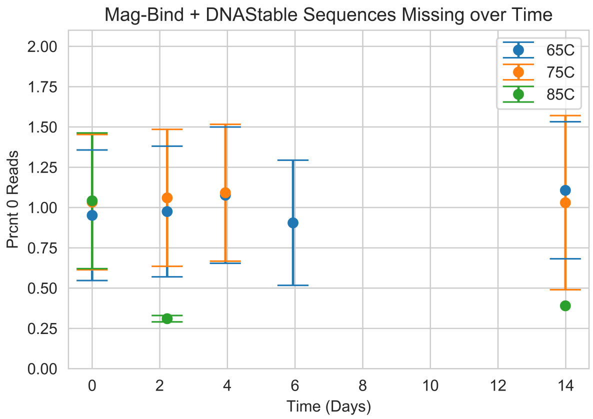

Supplementary Figure 11: Sequences missing from each time point and temperature for all triplicates, where error bars are the standard error from the mean.

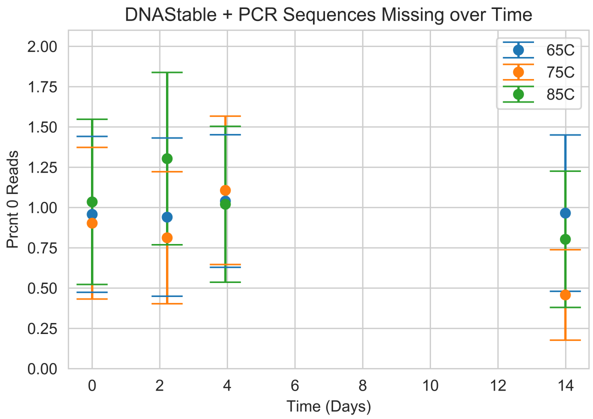

Supplementary Figure 12: Sequences missing from each time point and temperature for all triplicates, where error bars are the standard error from the mean.

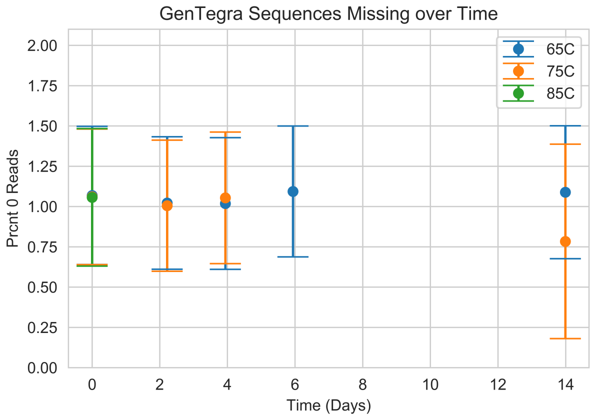

Supplementary Figure 13: Sequences missing from each time point and temperature for all triplicates, where error bars are the standard error from the mean.

##### 3.2 A sequence once lost is lost forever?

One might think a DNA sequence not seen by the sequencer in one time point is never recovered in all subsequent time points, but this is not the case. Typically, a sequence will be missing in one sample only to reappear later (as shown in the supplemental CSV files ending in ”missing sequences analysis”). This is likely due to the stochastic nature of sampling material to sequence. The fact that sequences tend not to stay missing supports the hypothesis that DNA degradation for these DNA sequences is stochastic.

For each temperature and time point, we present the data in three columns.

“Total percent missing” is calculated by finding the set of sequences that have zero coverage in ANY one of the triplicates and dividing that number by the number of total sequences (21,601 or 7,373, depending on the file being analyzed).

The “percent missing that reappear” examines all the sequences that are in the “total percent missing” set and determines what percent of those sequences are found either in another triplicate in the same time point, or a subsequent time point.

The “percent that stay missing” is calculated by finding the set of sequences that have zero coverage in all replicates in the examined time point as well as all subsequent time points, then dividing that number by the number of total sequences (21,601 or 7,373).

NaN (”not a number”) in the table means the file was not sequenced to sufficient sequencing depth at the given condition, had only one replicate, or the subsequent time point was not sequenced to sufficient sequencing depth and therefore we could not determine if the sequences stayed missing or not.

#### 3.3 Trimer analysis

In this analysis shown Supplementary Figures 14 and 15, we look at each trimer in each DNA sequence. For example, the sequence “ATCG” has the trimers “ATC” and “TCG”. In the last column, we also examine the sequence’s overall GC ratio.

We then look at each sample of DNA that was degraded and examine two groups of sequences: First, the sequences that were most sequenced (the top 5th percentile) and second, the sequences that were least sequenced (the bottom 5th percentile, including missing sequences).

We examined whether or not those two groups of sequences were statistically different with a t-test. By definition, this yielded many hundreds of p-values as there is one p-value for each trimer in each file in each method in each time point, temperature, and replicate.

Thus, we created a summary table in which we can clearly see that no trimer is statistically significantly different in all conditions, and the rates can vary quite a lot between the two files files aged in this experiment. The GC content metric is also not statistically significant in all conditions. Note that the primer sequences were not included in analysis here because they are constant for each file.

Even in instances where there is a statistically significant difference in the trimer presence after aging, for example, 92% for the trimer ”AGA” when looking at file 15 data, the number is quite different for file 8 where the trimer ”AGA” yields 74%. Or take the trimer ”AGT” with percentages of 62 for File 15 and 18 for File 8. This supports the hypothesis that there is no clear relationship between a particular trimer and increased degradation.

Percent of significant p-values over all temperatures and aging conditions by trimer - F8

| ACA | ACG | ACT | AGA | AGC | AGT | ATA | ATC | ATG | CAC | CAG | CAT |
|---|---|---|---|---|---|---|---|---|---|---|---|
| 92.0 | 18.0 | 14.0 | 92.0 | 47.0 | 18.0 | 91.0 | 8.0 | 2.0 | 35.0 | 70.0 | 50.0 |

| CGA | CGC | CGT | CTA | CTC | CTG | GAC | GAG | GAT | GCA | GCG | GCT |
|---|---|---|---|---|---|---|---|---|---|---|---|
| 14.0 | 39.0 | 42.0 | 4.0 | 88.0 | 92.0 | 21.0 | 80.0 | 33.0 | 9.0 | 21.0 | 85.0 |

| GTA | GTC | GTG | TAC | TAG | TAT | TCA | TCG | TCT | TGA | TGC | TGT |
|---|---|---|---|---|---|---|---|---|---|---|---|
| 32.0 | 57.0 | 64.0 | 40.0 | 73.0 | 16.0 | 1.0 | 78.0 | 78.0 | 0.0 | 87.0 | 70.0 |

| GC:AT |
|---|
| 85.0 |

Supplementary Figure 14: The percentage of significant t-test results comparing the most prevalent sequences to the least prevalent sequences across all conditions for file 8.

#### Percent of significant p-values over all temperatures and aging conditions by trimer – F15

| ACA | ACG | ACT | AGA | AGC | AGT | ATA | ATC | ATG | CAC | CAG | CAT |
|---|---|---|---|---|---|---|---|---|---|---|---|
| 81.0 | 65.0 | 2.0 | 74.0 | 13.0 | 62.0 | 83.0 | 4.0 | 55.0 | 67.0 | 67.0 | 27.0 |

| CGA | CGC | CGT | CTA | CTC | CTG | GAC | GAG | GAT | GCA | GCG | GCT |
|---|---|---|---|---|---|---|---|---|---|---|---|
| 6.0 | 6.0 | 60.0 | 4.0 | 74.0 | 72.0 | 11.0 | 18.0 | 65.0 | 72.0 | 13.0 | 79.0 |

| GTA | GTC | GTG | TAC | TAG | TAT | TCA | TCG | TCT | TGA | TGC | TGT |
|---|---|---|---|---|---|---|---|---|---|---|---|
| 60.0 | 72.0 | 79.0 | 30.0 | 69.0 | 11.0 | 0.0 | 81.0 | 74.0 | 0.0 | 74.0 | 81.0 |

| GC:AT |
|---|
| 81.0 |

Supplementary Figure 15: The percentage of significant t-test results comparing the most prevalent sequences to the least prevalent sequences across all conditions for file 15.

### 4 Overview Table

An comprehensive comparison of each preservation method. The times listed to encapsulate/dehydrate and de-encapsulate/re-hydrate serve to show the order of magnitude in which these processes currently take.

Note that Imagene does not list a half-life due to the fact that its samples degraded so little over time that no accurate half-life could be derived.

The DNAStable sample aged by ETH and the samples with no preservation other than dehydration degraded too quickly to measure adequately and so no accurate half-life could be derived.

| Method | Time to dehydrate/encapsulate | Half life per nt at 20C (years) | Order of time to rehydrate/de-encapsulate sample |
|---|---|---|---|
| Imagene | minutes | N/A | seconds |
| Trehalose | minutes | 4,900 | seconds |
| GenTegra | minutes | 3,000 | seconds |
| ETH Magnetic NP | minutes | 2,300 | minutes |
| Mag-Bind + DNAStable | minutes | 360 | minutes |
| Sugar Mix | minutes | 230 | seconds |
| ETH Trehalose | minutes | 170 | seconds |
| Filter Paper | minutes | 100 | seconds |
| DNAStable + PCR | minutes | 45 | seconds |
| DNAStable | minutes | 19 | seconds |
| ETH DNAStable | minutes | N/A | seconds |
| No Additives | N/A | N/A | seconds |

Supplementary Table 4: An overview of the storage methods explored.

### 5 Ligation Protocol

The ligation protocol used is a combination of the Illumina TruSeq ChIP Sample Preparation protocol and the Illumina TruSeq Nano Library Prep protocol, using the Illumina TruSeq Nano reagents. The following are the step-by-step directions used in this work.

1. Add 40 $\mu$L ERP2 to each well with 60 $\mu$L of PCR product

2. Pipette up and down to mix

3. Place on the thermal cycler and run the ERP program (30 min at 30C) then place on ice. Each well contains 100 $\mu$L.

4. Add 160 $\mu$L well-mixed AMPure XP Beads to each well of the PCR plate containing 100 $\mu$L End Repair Mix.

5. Gently pipette the entire volume up and down 10 times to mix thoroughly.

6. Incubate the PCR plate at room temperature for 15 minutes.

7. Place the PCR plate on a magnetic stand at room temperature for 15 minutes or until the liquid is clear.

8. Using a 200 $\mu$L single channel or multichannel pipette set to 127.5 $\mu$L, remove and discard 127.5 $\mu$L of the supernatant from each well of the PCR plate.

9. Repeat step 8 one time.

   NOTE- Leave the PCR plate on the magnetic stand while performing the following 80% EtOH wash steps (10-12).

10. With the PCR plate on the magnetic stand, add 200 $\mu$L freshly prepared 80% EtOH to each well without disturbing the beads.

11. Incubate the PCR plate at room temperature for 30 seconds, and then remove and discard all of the supernatant from each well. Take care not to disturb the beads.

12. Repeat steps 10 and 11 one time for a total of two 80% EtOH washes.

13. Let the PCR plate stand at room temperature for 15 minutes to dry, and then remove the plate from the magnetic stand.

14. Resuspend the dried pellet in each well with 17.5 $\mu$L Resuspension Buffer (RSB). Gently pipette the entire volume up and down 10 times to mix thoroughly.

15. Incubate the PCR plate at room temperature for 2 minutes.

16. Place the PCR plate on the magnetic stand at room temperature for 5 minutes or until the liquid is clear.
17. Transfer 15 $\mu$L of the clear supernatant from each well of the PCR plate to the corresponding well of a new 96-well 0.3 ml PCR plate.
18. Add 15 $\mu$L of A Tailing Ligase (ATL) to the 15 $\mu$L of supernatant from previous step.
19. Briefly spin down on a standard, small benchtop centrifuge.
20. Place on the thermal cycler and run the ATAIL70 program listed below. Each well contains 30 $\mu$L.
    (a) Choose the preheat lid option and set to 100C
        i. 37C for 30 minutes
        ii. 70C for 5 minutes
        iii. 4C for 5 minutes
21. Add the following reagents from the Illumina TruSeq Nano ligation kit IN ORDER:
    (a) RSB (2.5 $\mu$L)
    (b) LIG2 (2.5 $\mu$L)
    (c) DNA adapter (2.5 $\mu$L)
22. Pipette up and down, centrifuge briefly
23. Run lig program listed here in step a.
    (a) Choose the preheat lid option and set to 100C
        i. 30C for 10 minutes
        ii. Hold at 4C
    (b) Add 5 $\mu$L STL to each well, and then mix w/pipette. NOTE- You can hold this mixture at -20C overnight with no trouble.
24. Vortex Sample Purification Beads (SPB) until well dispersed.
25. Perform steps a through l using the Round 1 volumes.
    (a) Add SPB to each well, and then mix thoroughly as follows.
        i.

| | |
|---|---|
| Round 1 | 42.5 $\mu$L |
| Round 2 | 50 $\mu$L |

    (b) Pipette up and down.
    (c) Incubate at room temperature for 5 minutes.
    (d) Place on a magnetic stand and wait until the liquid is clear (2–5 minutes).

(e) Remove and discard all supernatant from each well.

(f) Wash 2 times as follows.

  i. Add 200 $\mu$L freshly prepared 80% EtOH to each well.
  ii. Incubate on the magnetic stand for 30 seconds.
  iii. Remove and discard all supernatant from each well.

(g) Use a 20 $\mu$L pipette to remove residual EtOH from each well.

(h) Air-dry on the magnetic stand for 5 minutes.

(i) Add RSB to each well.

  i.

| | |
|---|---|
| Round 1 | 52.5 $\mu$L |
| Round 2 | 27.5 $\mu$L |

(j) Remove from the magnetic stand, and then mix thoroughly as follows. Pipette up and down.

(k) Incubate at room temperature for 2 minutes.

(l) Place on a magnetic stand and wait until the liquid is clear (2–5 minutes).

(m) CONTINUE TO STEP 26

26. ROUND1- Transfer 50 $\mu$L supernatant to the corresponding well of the CAP plate.

27. Repeat steps a through l with the new plate using the Round 2 volumes.

28. ROUND2- Transfer 25 $\mu$L supernatant to the corresponding well of the PCR plate.

SAFE STOPPING POINT If you are stopping, seal the plate and store at -25C to -15C for up to 7 days.

At this point, ligation and purification is done.

The following is the recipe for each ligated sample:

| Sample | Concentration | Volume ($\mu$L) |
|---|---|---|
| DNA mix | | 3 |
| Forward post-ligation primer (see **Supplemental Section 1**) | 10˘M | 1.5 |
| Reverse post-ligation primer (see **Supplemental Section 1**) | 10˘M | 1.5 |
| Kapa HiFi 2x | 2.5x | 12 |
| Molecular Grade Water | | 12 |
| Total | | 30 |

Follow the following thermocycling protocol with the above 30 $\mu$L mixture:

[noitemsep]Choose the preheat lid option and set to 100C 95C for 3 minutes 8 total cycles of:[noitemsep]

- – 98C for 20 seconds

  - 60C for 15 seconds
  - 72C for 30 seconds
- 72C for 3 min
- Hold at 4C

#### 6 Arrhenius Equation

Assuming DNA degradation is a first-order decay process, then the rate of the reaction is dependent on the rate constant, k, and the concentration of the DNA, $[DNA]$.

$$r = k[DNA] \tag{1}$$

From equation (1), we can derive the relationship between k and the initial and end concentrations of DNA for a given time, t.

$$ln[DNA] = -kt + ln[DNA]_0 \tag{2}$$

The temperature dependence of k can be modeled by the Arrhenius equation, where A is the pre-exponential factor, Ea is the activation energy, R is the gas constant, and T is temperature.

$$k = Ae^{\frac{-E_a}{RT}} \tag{3}$$

Taking the natural logarithm of equation (3), we can calculate Ea.

$$ln[k] = \frac{-E_a}{RT} + ln[A] \tag{4}$$

The half-life of the DNA ($t_{1/2}$) can be calculated by solving equation (2), where at $t_{1/2}$ the concentration of DNA is half of the original concentration.

$$t_{(1/2)} = \frac{ln(2)}{k} \tag{5}$$
